# Loss of C3aR induces immune infiltration and inflammatory microbiota in a new spontaneous model of colon cancer

**DOI:** 10.1101/2021.01.18.426963

**Authors:** Carsten Krieg, Lukas M. Weber, Bruno Fosso, Gary Hardiman, Erika Mileti, Sahar El Aidy, Marinella Marzano, Mark D. Robinson, Silvia Guglietta

## Abstract

Several lines of evidence suggest that inflammation plays a pivotal role in the development and progression of colorectal cancer (CRC) and can be unleashed by the loss of innate immunosurveillance. The complement system is a well characterized first line of defense against pathogens and a central component of the immune response. Emerging evidence suggests that complement anaphylatoxin C3a produced upon complement activation and acting via its receptor (C3aR) may play a role in intestinal homeostasis. However, to date, it is unknown whether and how the C3a/C3aR axis can affect CRC. By mining publicly available datasets, we found that CpG island methylation of *c3ar1* occurs in CRC patients and is associated with significant downregulation of C3aR. By reverse-translating this finding we were able to shift in APC^Min/+^ mice the tumorigenesis from the small intestine to the colon therefore generating a novel mouse model, which more closely mirrors the CRC in humans. Transcriptomic analysis on colorectal polyps from our newly developed genetic mouse model revealed a significant increase in innate and adaptive immune signatures in absence of C3aR. Furthermore, loss of C3aR significantly impacted the fecal and tumor-associated microbiota and supported the blooming of pro-inflammatory bacterial species as confirmed by experiments of fecal microbiota transplantation.

Future studies will elucidate whether loss of C3aR can be exploited as a biomarker for sub-groups of CRC and whether the C3a/C3aR axis may be exploited for the generation of more effective therapeutic interventions.

## INTRODUCTION

Colorectal cancer (CRC) is the third most diagnosed cancer worldwide and the first cause of non-smoking-related cancer deaths and it is expected to increase by 60% with 2.2 million new cases and 1.1 million deaths annually by the year 2030 (Arnold et al., 2017; Siegel et al., 2018). This trend is even more alarming considering that recent data show increased incidence of CRC in young adults, who often experience more aggressive disease and lower survival than the older population (Siegel et al., 2020). While early screening, surgery and adjuvant therapies have significantly improved CRC outcome, still about 30% of patients with CRC undergo recurrence and develops metastatic disease, therefore suggesting that the discovery of new treatments with improved efficacy is highly desirable. A major obstacle to the identification of more effective therapies is the lack of suitable preclinical mouse models that closely mirror the multi-step process of CRC development in humans. Attempts to model human CRC using transgenic mice carrying known genetic mutations that occur in human CRC resulted in the development of several models, which share similar limitations: lack of invasive phenotype, development of multiple adenomas localized in the small intestine rather than in the colon, low penetrance and low reproducibility (Burtin et al., 2020; Fodde et al., 2001). This is not surprising if we consider that genetic mutations represent only a piece of the very complex puzzle of human CRC, where epigenetic alterations, environmental and host-associated factors are emerging as major determinants in the origin and the heterogeneity of the tumors (Pancione et al., 2012). Therefore, identifying additional events that primarily cause tumor initiation may represent a successful strategy to more closely model the human disease in mice. The gastrointestinal tract is among the largest mucosal surfaces in the body and mounting prompt and efficacious innate immune responses is indispensable to maintaining homeostasis and preventing harmful inflammation. In support of this observation, deficiencies in innate sensing mechanisms have been associated with increased intestinal inflammation and CRC in human and mouse models (Coleman et al., 2018; de Souza et al., 2018; Garrett et al., 2007; Rakoff-Nahoum and Medzhitov, 2007; Salcedo et al., 2010). The complement system represents the first line of defense against pathogens and is a central player in immunity due to its ability to establish extensive networks with other innate immune pathways (Arbore et al., 2016; Hajishengallis and Lambris, 2010; Ostvik et al., 2014; Zhang et al., 2007).

We previously reported that activation of the alternative complement pathway occurs during spontaneous small intestinal tumorigenesis and that the C3a-C3aR axis is involved in tumor-associated thrombosis via promotion of NET formation (Guglietta et al., 2016). Furthermore, in a model of intestinal ischemia/reperfusion injury, the C3a-C3aR axis proved essential for tissue regeneration and protection from oxidative damage (Wu et al., 2013). In light of these findings, which suggest a critical function for the C3a/C3aR axis in intestinal homeostasis, we set out to investigate its role in the development of human CRC. By mining publicly available gene expression datasets, we found that *c3ar1* in CRC patients is among the top 1% down-regulated genes. Notably, we found that *c3ar1* down-regulation in human is primarily associated with CpG island methylation rather than somatic mutations. We then reverse translated our finding from CRC patients into APC^Min/+^ mice, which carry a mutation in the *apc* gene and mainly develop polyps in the small intestine (Fodde et al., 2001; Guglietta et al., 2016). Notably, by knocking down C3aR, therefore generating APC^Min/+^/C3aR-/- mice, we observed an unprecedent shift of tumor growth from the small intestine to the colon and a significant induction of innate and adaptive immune responses. These events were dependent on gut microbiota. Indeed, the disruption of the C3a/C3aR axis affected both the fecal and tumor-associated microbiota and fecal microbiota transplantation effectively reproduced in APC^Min/+^ mice the immune infiltrate and tumor growth observed in APC^Min/+^/C3aR-/- mice. Altogether, based on our findings in human CRC, by disrupting the C3a/C3aR axis, we generated a new spontaneous CRC models whereby immune responses, microbiota and genetic alterations contribute to the development of tumors in the colon, more closely resembling the human disease. Future studies will determine whether the absence of C3aR could be exploited for therapeutic intervention and whether C3aR could serve as biomarker for diagnostic and therapeutic purposes in CRC.

## RESULTS

### *c3ar1* down-regulation occurs in CRC patients

Analysis of the gene expression libraries provided through the Skrzypczack Colorectal cohort, Sabates-Bellver Colon cohort, Ki Colon cohort and TCGA cohort (https://www.oncomine.org), showed that in sub-groups of both rectal and colon cancer patients *c3ar1* is downregulated compared to controls (Fig. 1a). Because mutations of single genes in the complement pathway are very rare, we reasoned that down-regulation of the *c3ar1* gene could be more efficiently mediated by epigenetic modifications (Olcina et al., 2018). Methylation occurs at high frequency in CRC and it has been shown to also induce silencing of tumor suppressor genes (Tse et al., 2017). In order to assess this possibility, we mined the MethHC database of DNA methylation and gene expression in human cancer and found that *c3ar1* gene is more highly methylated in CRC tissues as compared to the paired normal mucosa (Huang et al., 2015). Remarkably, significant hypermethylation was seen in the N and S shores (regions up to 2 kb away from CpG islands) and in the shelves (Fig. 1b-d). As shown by Irizarry and collaborators, DNA methylation in the N and S shores of CpG islands inversely correlates with gene expression levels (Irizarry et al., 2009). These results suggest that the downregulation of C3aR in CRC could be due to CpG island methylation events, that have been shown to be prominent for the development of CRC (Jung et al., 2020). Taken together these findings support the hypothesis that C3aR plays a previously unrecognized role in intestinal homeostasis and loss or reduction of its expression due to DNA methylation or mutations occurs in CRC patients.

**Figure 1.**
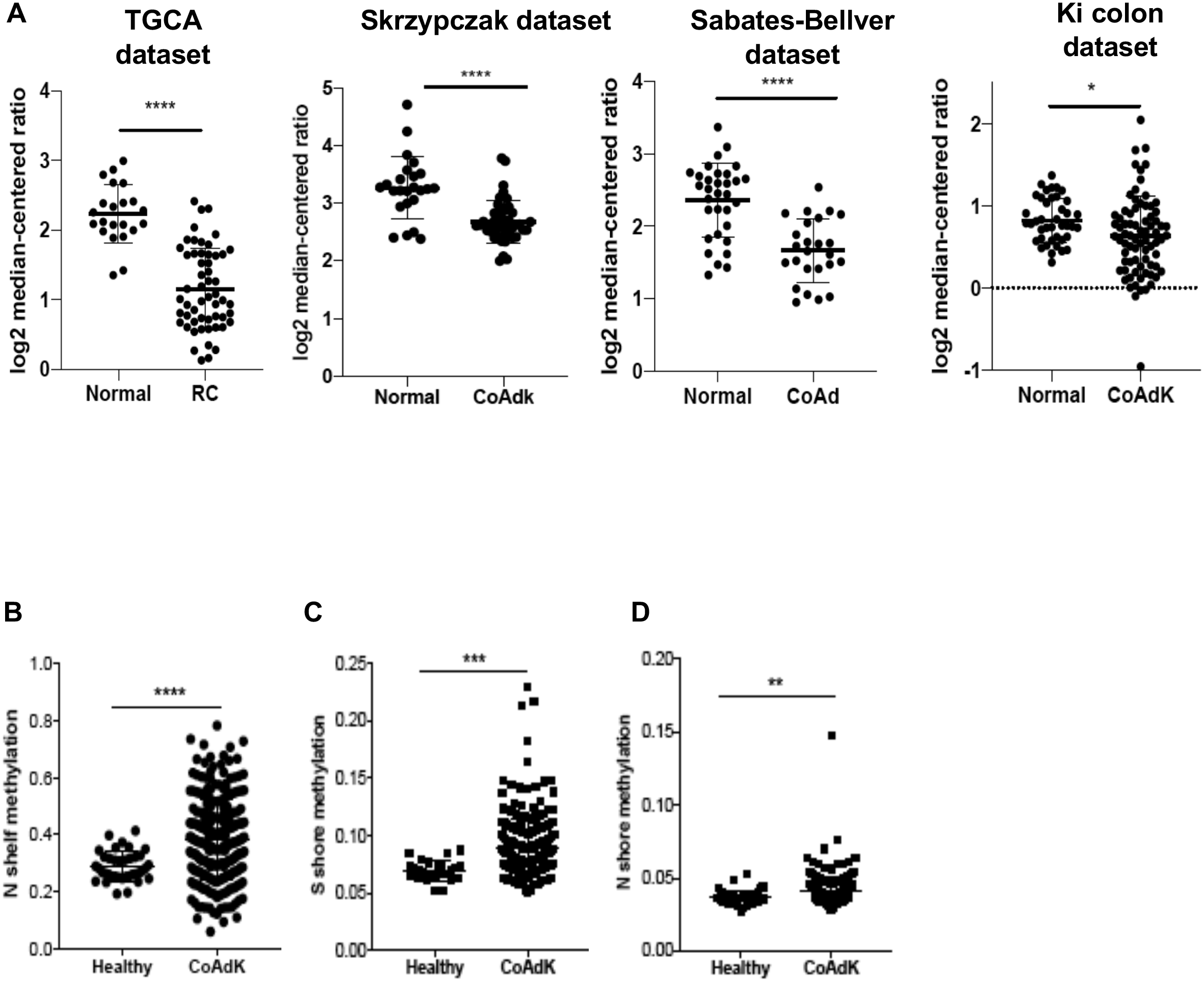
C3aR methylation and down-regulation in CRC patients. (A) C3aR expression in patients with rectal and colon cancer from 4 independent datasets (TGCA: 22 healthy-55 RC; Skrzypczak cohort: 24 healthy-45CoAdK; Sebates-Bellver cohort:32 healthy-25 CoAd; Ki cohort: 41 healthy-76 CoAdK). (B) N shelf, (C) S shore and (D) N shore methylation of *c3ar1* methylation in patients with CRC. Significance was calculated using t-test (*p>0.05; **** p>0.0001).

### C3a/C3aR axis plays a previously unappreciated role in CRC development

While we found that C3aR is downregulated in CRC patients, the functional significance of this phenomenon for the tumorigenic process is unknown. To test the role of the C3a/C3aR axis in CRC development, we reverse translated our finding from the database into mouse models. First, age-and sex-matched C3aR-/- and WT mice were administered the carcinogen azoxymethane (AOM) followed by three cycles of the inflammatory agent dextran sodium sulphate (DSS) and their body weight was monitored every other day until day 70, when tumor number and size were quantified. Relative to WT, C3aR-/- mice showed increased weight loss and reduced survival throughout the experiment (Fig. 2a-b) suggesting that C3aR protects from excessive intestinal inflammation. This result is in agreement with previously published literature showing that lack of complement activation in C3-/-mice exacerbates chronic intestinal inflammation (Elvington et al., 2015). Notably, as shown in Fig. 2c-e, at the end of the experiment, C3aR-/- mice developed significantly higher tumor number and higher tumor load in the colon as compared to the WT mice, with predominance of larger tumors (> 2mm). As innate and adaptive immune responses are both involved in colon inflammation and tumorigenesis, we set up three interlinked flow cytometry panels to characterize the immune infiltrate in the tumors and in the mesenteric lymph nodes (mLN). As shown in Fig. 2f-h, we found higher numbers of CD4+ IL17A+ (Th17) and IFN-*γ*+ T cells (Th1) cells and increased CD11c+ macrophages in the mLN of C3aR-/- mice with no significant differences in the amount of FoxP3+CD25+ Tregs. In contrast, while no differences relative to WT were observed in dendritic cell and macrophage subsets in the tumors of C3aR-/- mice (Fig. 2i), we found significantly higher numbers of Tregs, and IFN-*γ*+/IL17A+ CD4+T cells (Th1/Th17) compared to WT mice (Fig. 2j-k). Although more than 20% of individuals with IBD develop CRC, intestinal inflammation accounts for only about 2% of total CRC (Munkholm, 2003). Therefore, we next investigated the impact of C3aR loss in the APC^Min/+^ spontaneous model, which like 80% of human CRC (Polakis, 2012), carries a mutation of the *apc* gene. We generated APC^Min/+^ mice lacking the *c3ar1* gene (APC^Min/+^/C3aR-/-) and compared tumor development in the colon from 5 to 28 weeks of age in the APC^Min/+^ and APC^Min/+^/C3aR-/- mice. We and others have shown that in APC^Min/+^ mice CRC development is sporadic, with the highest tumor burden in the distal small intestine (Fodde et al., 2001; Guglietta et al., 2016; Haigis et al., 2004). In line with the human data and the results in the model of inflammation-driven CRC, we found that in the absence of C3aR, starting at 10 weeks of age, APC^Min/+^ mice showed a shift of tumorigenesis from the small intestine to the colon with the highest localization in distal colon and rectum (Fig. 3a and b). Similar to the approach used in the AOM/DSS model, we performed flow cytometry to quantify the immune infiltrate associated with tumor development. Due to the very low number of colon tumors in APC^Min/+^ mice, we analyzed the mLN and the colon lamina propria (cLP) by flow cytometry. Similar to the results obtained in the AOM/DSS model, loss of C3aR resulted in significantly higher numbers of Th17, Th1, Th1/Th17 and CD8+ T cells in the cLP of APC^Min/+^/C3aR-/- mice compared to the APC^Min/+^ counterpart (Fig. 3c-h). In the mLN we found higher numbers of CD4+ and CD8+T cells in the APC^Min/+^/C3aR-/- mice compared to APC^Min/+^ mice. However, in terms of functional differences, we only observed increased number of IFN-*γ*-producing CD8+T (Suppl. Fig. 1a-f). The role and prognostic significance of infiltrating immune cells in the lamina propria and tumors of C3aR-/- mice is currently unknown. However, the coexistence of T regs, Th17 and IFN-*γ*-producing CD8+T cells in the tumors and cLP of both mouse models is in line with published literature showing that Tregs can promote Th17 cell infiltration during CRC and that Th17 in human CRC recruit CD8+T cells (Amicarella et al., 2017; Watanabe et al., 2016). Altogether by disrupting the C3a/C3aR axis, we generated a novel mouse model, which spontaneously develops colon tumors and therefore more closely mirrors the disease in humans. Our findings confirm the data in CRC patients and suggest that the C3a/C3aR axis play a previously unappreciated role in colon cancer and inflammation.

**Figure 2.**
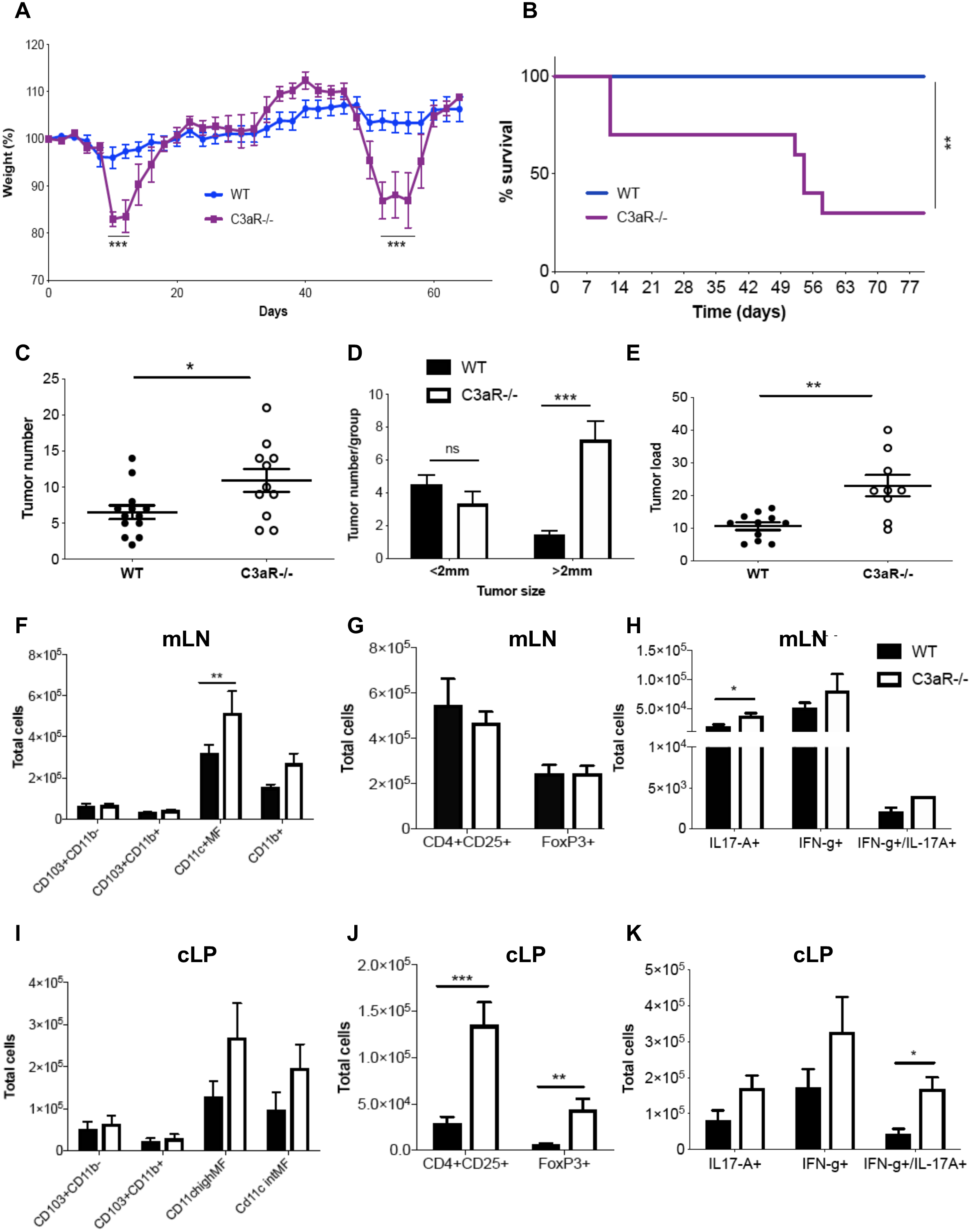
Loss of C3aR exacerbate inflammation and tumor development in inflammation-driven CRC. (A) WT and C3aR-/- mice were treated with AOM/DSS and weight loss was monitored every other day. (B) Overall survival in AOM/DSS-treated WT and C3aR-/- mice. (C) Total number of tumors in the colon of WT and C3aR-/- mice. (D) Single tumor diameters in WT and C3aR-/- mice were measured with a sliding caliper and assigned to the groups <2MM or >2mm. (E) Tumor load for each mouse was calculated by adding up individual tumor diameters. Single cell suspensions from mesenteric lymph nodes (mLN) and tumors of WT and C3aR-/- mice were analyzed by flow cytometry. Display of total numbers of (F) myeloid cells (CD11c MF = CD11c+ macrophages), (G) Tregs (CD3+ CD4+ CD25+ FoxP3+), and (H) Th1 (CD3+CD4+ IFN*γ*+) cells, Th17 (CD3+CD4+ IL-17A+) cells and Th1/Th17 (CD3+CD4+ IFN-*γ*+IL-17A+) cells in mLN. Total numbers of (I) myeloid cells, (K) Tregs, and (J) Th1, Th17 and Th1/Th17 in tumors as gated in F-H. Results are pooled from two independent experiments with a minimum of 8 mice/group). Significance was calculated in A, C and E using t-test and in B using Mantel-Cox test; 2-way ANOVA with Bonferroni post-test was used in panel D and F-L (* p< 0.05; ** p< 0.01; *** p< 0.001).

**Figure 3.**
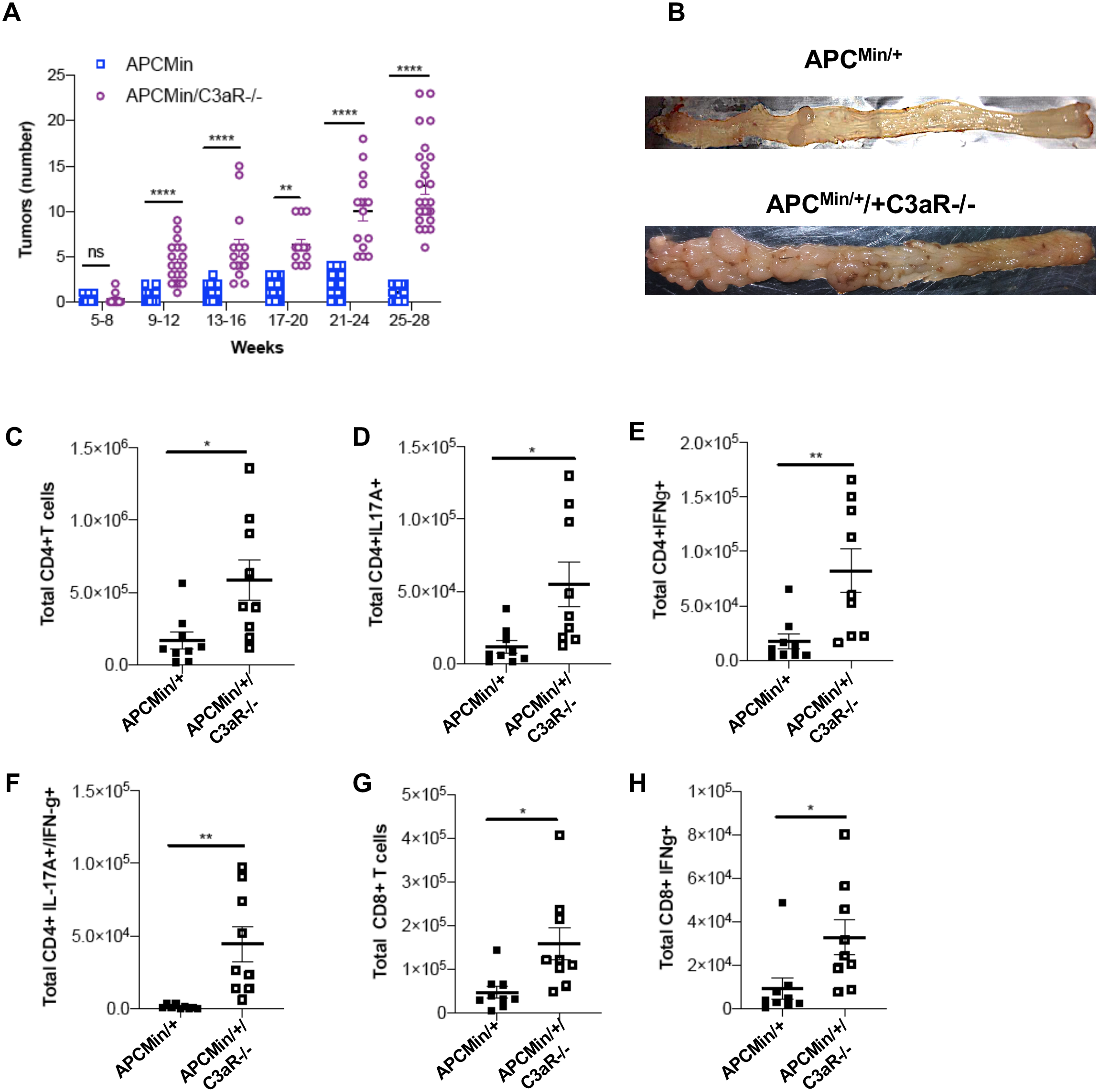
Loss of C3aR switches tumorigenesis from small intestine to colon in APCMin/+ and mice and promotes increased infiltration with Th1/Th17. (A) Tumor number in the colon of APC^Min/+^ and APC^Min/+^/C3aR-/- mice was assessed starting at 5 weeks until 28 weeks of age. (B) Display of an example of tumor number and distribution in the colon of APC^Min/+^ and APC^Min/+^/C3aR-/- mice. Single cell suspensions from colon lamina propria (cLP) APC^Min/+^ and APC^Min/+^/C3aR-/- mice were analyzed by FACS and total number of (C) CD4+T cells, (D) Th17 cells, (E) Th1 cells, (F) Th1/Th17 cells, (G), CD8+ T cells (H) Tc cells (CD3+CD8+IFN-*γ*+) were gated and calculated as for Fig. 2. In panel A significance was calculated by using 2-way ANOVA with Bonferroni post-test (* p< 0.05; ***p< 0.001) and a minimum of 10 mice/group was used. In panels C-H a minimum of 7 animals/group was used and significance was calculated using unpaired t-test (ns= not significant; * p< 0.05; ** p< 0.01).

### Disruption of the C3a/C3aR axis results in transcriptional up-regulation of innate and adaptive immune pathways in the healthy distal colon of APC^Min/+^/C3aR-/- mice

To understand how lack of C3aR shapes the tumor microenvironment and the adjacent healthy mucosa in our newly developed spontaneous mouse model of colon tumorigenesis, we performed RNA-Seq analysis in 8 and 12-week-old APC^Min/+^ and APC^Min/+^/C3aR-/- and used WT and C3aR-/-littermates as respective controls. Noting that absence of C3aR induced prevalent development of tumors in the distal colon and rectum, we suspected that the absence of C3aR might have a more profound effect on the microenvironment of the distal colon. Therefore, for RNA-Seq studies, proximal and distal colon were kept separated. When performing the comparison between the healthy colons among all the strains and the subsequent gene clustering to generate the heatmap, we observed that APC^Min/+^ mice had a more similar transcriptomic profile to WT mice than to APC^Min/+^/C3aR-/- mice, and similarly, healthy colons of C3aR-/- mice exhibited a transcriptomic profile more similar to the colons of APC^Min/+^/C3aR-/-than to the colons of WT mice (Fig. 4a and Suppl. Fig. 2a-f). These results show that the absence of C3aR, rather than the presence of tumors, shapes the transcriptomic profile of tumor-adjacent healthy mucosa. Using the edgeR-GLM framework (see Methods), we then tested the differential gene expression profile in distal colon and tumors of APC^Min/+^ compared to APC^Min/+^/C3aR-/- mice. As shown in Suppl. Fig. 2f, by using a 2-fold threshold, C3aR loss resulted in a significant change in the expression level of 318 genes in the healthy distal colon, out of which 235 were up-regulated and 83 down-regulated. Notably, iPathway analysis revealed that the top 25 Gene Ontology (GO) biological processes associated with differentially expressed genes were related to innate and adaptive immune responses (Fig. 4b). Accordingly, in the distal colon of APC^Min/+^C3aR-/- mice, we observed a significant up-regulation of genes involved in the response to bacteria (*nos2, nox1, ltf* and *s100a9);* adaptive immune responses-related genes (*il18bp*, *ifn-γ*, *granzyme a* and *b, klrc3),* ifn-*γ*-mediated signaling genes *(stat1, nlrc5, gbp2).* In contrast, in the distal colon of APC^Min/+^/C3aR-/- mice we found a significant down regulation of defensin genes involved in the response to bacteria such as *reg1*and *reg3a,* (Cash et al., 2006) and several genes encoding for solute carrier transporters such as *slc30a10*, *slc35e3, slc20a1* and *slc10a2* (Fig. 4c).

**Figure 4.**
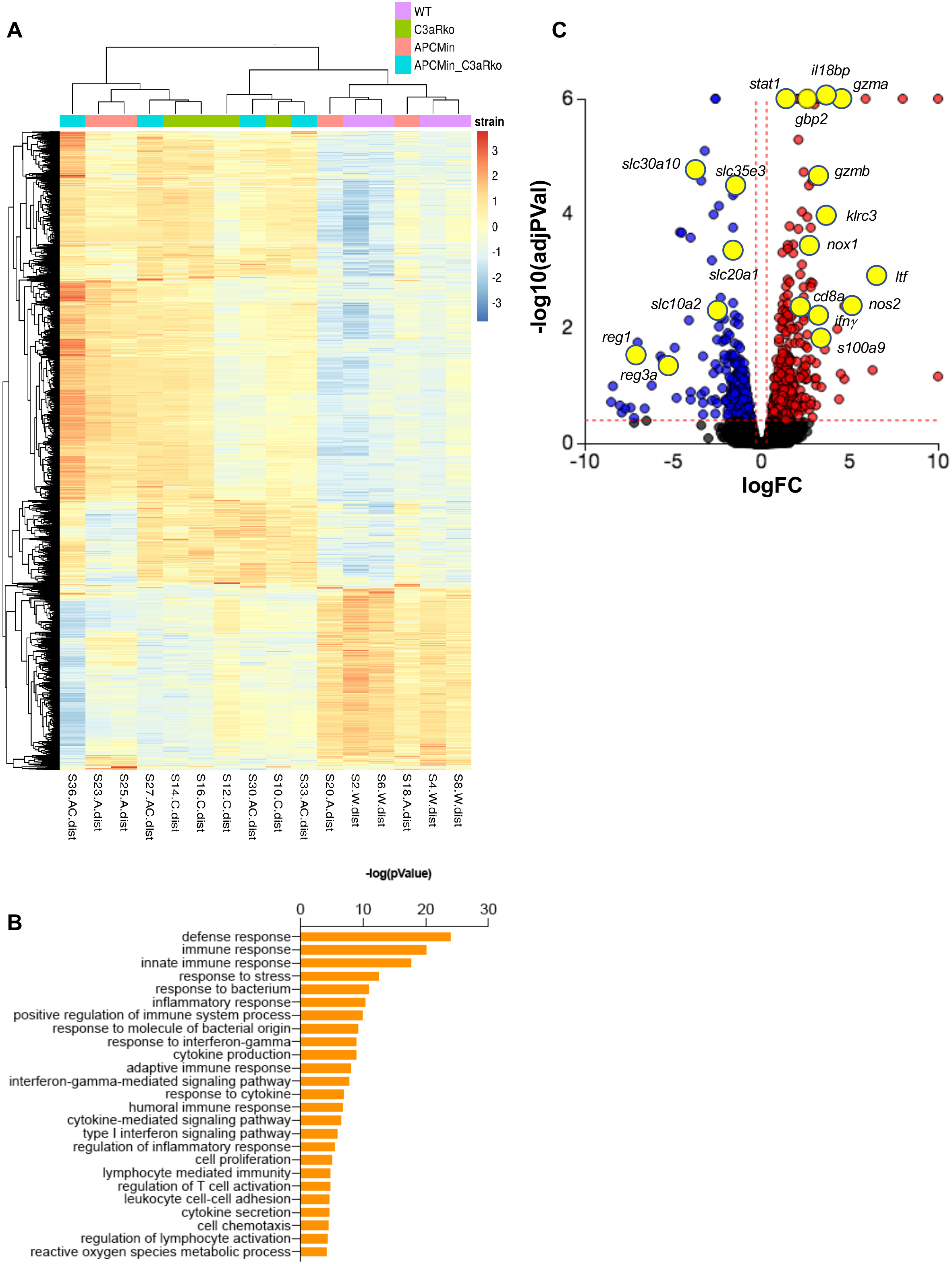
Loss of C3aR results in transcriptional up-regulation of innate and adaptive immune pathways in the healthy distal colon of APC^Min/+^/C3aR-/- mice. (A) Heatmap from (row-wise z-transformed) log counts per million values, using data from distal colon of indicated mouse strains and all genes that are significant at 5% FDR in at least one contrast. Rows represent genes, and columns represent individual samples. Each row (gene) is z-transformed to have mean zero and standard deviation one. The heatmap also includes column annotation labels indicating the mouse strain for each sample. (B) Visualization of the gene-associated Gene Ontology (GO) biological processes in healthy colon. (C) Volcano plot illustrating the magnitude of fold change for all genes differentially expressed in the distal colon of APC^Min/+^C3aR-/-vs APC^Min/+^ mice (n=4). Representative significantly up-regulated and down-regulated genes belonging to the GO biological processes shown in B are highlighted in yellow. Significance was calculated using the edgeR function decideTestsDGE with Benjamini-Hochberg correction for false discovery rate (FDR); a default FDR threshold of 0.05 and a log2 fold change (log2FC) threshold of 0.6 were applied.

### Colon tumors from APC^Min/+^/C3aR-/-show an enrichment in inflammatory immune pathways

We next compared the transcriptomic profile of colon tumors developed by APC^Min/+^ and APC^Min/+^/C3aR-/- mice. Using a 2-fold threshold, we found 72 downregulated genes and 369 upregulated genes in colon polyps from APC^Min/+^/C3aR-/- mice, clearly indicating that loss of C3aR affects the transcriptomic profile of the polyps (Fig. 5a).

**Figure 5.**
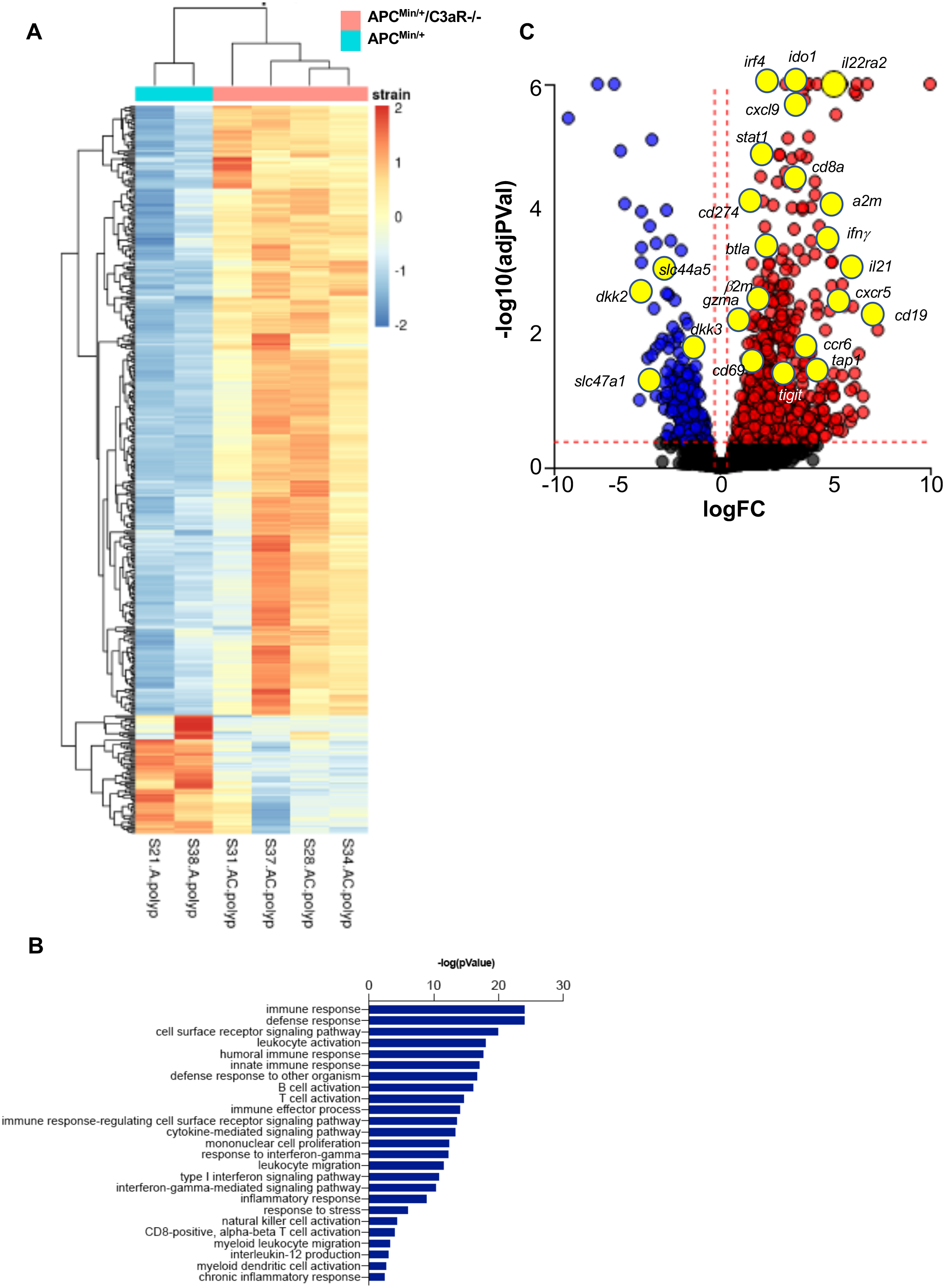
APC^Min/+^/C3aR-/- polyps show an enrichment of inflammatory pathways. (A) Heatmap from log counts per million values using data from colon polyps from APC^Min/+^ and APC^Min/+^/C3aR-/- mice. (B) Visualization of the gene-associated Gene Ontology (GO) biological processes in the polyps. (C) Volcano plot illustrating the magnitude of fold change for all genes differentially expressed in the polyps of APC^Min/+^C3aR-/- vs APC^Min/+^ mice (n=4). Representative significantly up-regulated and down-regulated genes belonging to the GO biological processes shown in B are highlighted in yellow. Vertical and horizontal red dotted lines indicate the threshold. Significance was calculated using the edgeR function decideTestsDGE with Benjamini-Hochberg correction for false discovery rate (FDR); a default FDR threshold of 0.05 and a log2 fold change (log2FC) threshold of 0.6 were applied.

In line with the results from the healthy mucosa of the distal colon, iPathway analysis revealed that the top 25 Gene Ontology (GO) biological processes associated with differentially expressed genes in the polyps were related to innate and adaptive immune responses (Fig. 5b). Accordingly, in the tumors of APC^Min/+^/C3aR-/- mice, we found significant up-regulation of genes associated with defense response, such as *cd8, ccr6, il21, cd19, il22ra2;* IFN-*γ−*mediated signaling pathway-related genes, such as *ifi* and guanylate-binding protein (GBP) family genes *tap-1, ciita, β2-microglobulin* and *stat1*; genes associated with development and maturation of innate immune cells and response to type I interferon such as *irf4* and *irf8* and *cd83*; negative regulator of the immune response, such as *btla, cd274 (pd-l1), cd40lg* and *ido1*; genes associated with leucocyte migration such as, *cxcl9, cxcl10, cxcl11* and *cxcr5*; NK cell activation genes such as *il-12b* and *ncr1*. Among the significantly down-regulated genes, we found genes for transporters including *slc47* and *slc44a5* and Wnt signaling pathway inhibitors such as *dkk2* and *dkk3* (Fig. 5c). These latter have recently been shown to be up-regulated following *apc* loss in CRC patients and to mediate the failure of anti-PD1 treatment in MSS-high CRC by dampening NK and CD8-mediated anti-tumor immune responses (Xiao et al., 2018). Altogether, the results from our transcriptomic analysis suggest that loss of C3aR promotes a vigorous inflammatory signature in the colon and in the tumors of APC^Min/+^ mice.

### APC^Min/+^/C3aR-/- microbiota transplantation transfers tumor growth to APC^Min/+^ mice

Because evidence suggests that CRC tumor development is strictly influenced by the microbial flora (Ahn et al., 2013; Arthur et al., 2014; Dejea et al., 2018; Tilg et al., 2018; Wu et al., 2009), we sought to understand whether the transfer of APC^Min/+^/C3aR-/- microbiota could increase tumor development in the colon of APC^Min/+^ mice. Tumor-free, five-week-old APC^Min/+^ mice were treated with broad-spectrum antibiotics, then administered via gavage with fecal microbiota from 12- week-old APC^Min/+^/C3aR-/- mice or APC^Min/+^ mice and sacrificed at the age of 12 weeks (Fig. 6a). As shown in Fig. 6b, while APC^Min/+^ mice receiving the APC^Min/+^ microbiota developed very few tumors in the colon, the transfer of APC^Min/+^/C3aR-/- microbiota resulted in increased tumor burden in the colon, with no changes in the total number of small intestinal tumors (Suppl. Fig. 3a-b). To test whether the microbiota could impact immune cell recruitment in the colon of microbiota-transplanted mice, we used flow cytometry to characterize the immune cell infiltrate in the APC^Min/+^ mice receiving APC^Min/+^ or APC^Min/+^/C3aR-/- microbiota. As shown in Fig. 6c-h and Suppl. Fig. 3c-h, APC^Min/+^ mice receiving APC^Min/+^/C3aR-/- feces showed higher tumor numbers in the colon and increased Th1, Th17 and Th1/Th17 cell infiltration in the lamina propria. Altogether, these data clearly demonstrated that, besides affecting colon tumor development, the microbiota transplantation from APC^Min/+^/C3aR-/-, but not APC^Min/+^ mice, recapitulated the immune signature that we described at protein and RNA level in colon and tumors of mice lacking the C3aR signaling.

**Figure 6.**
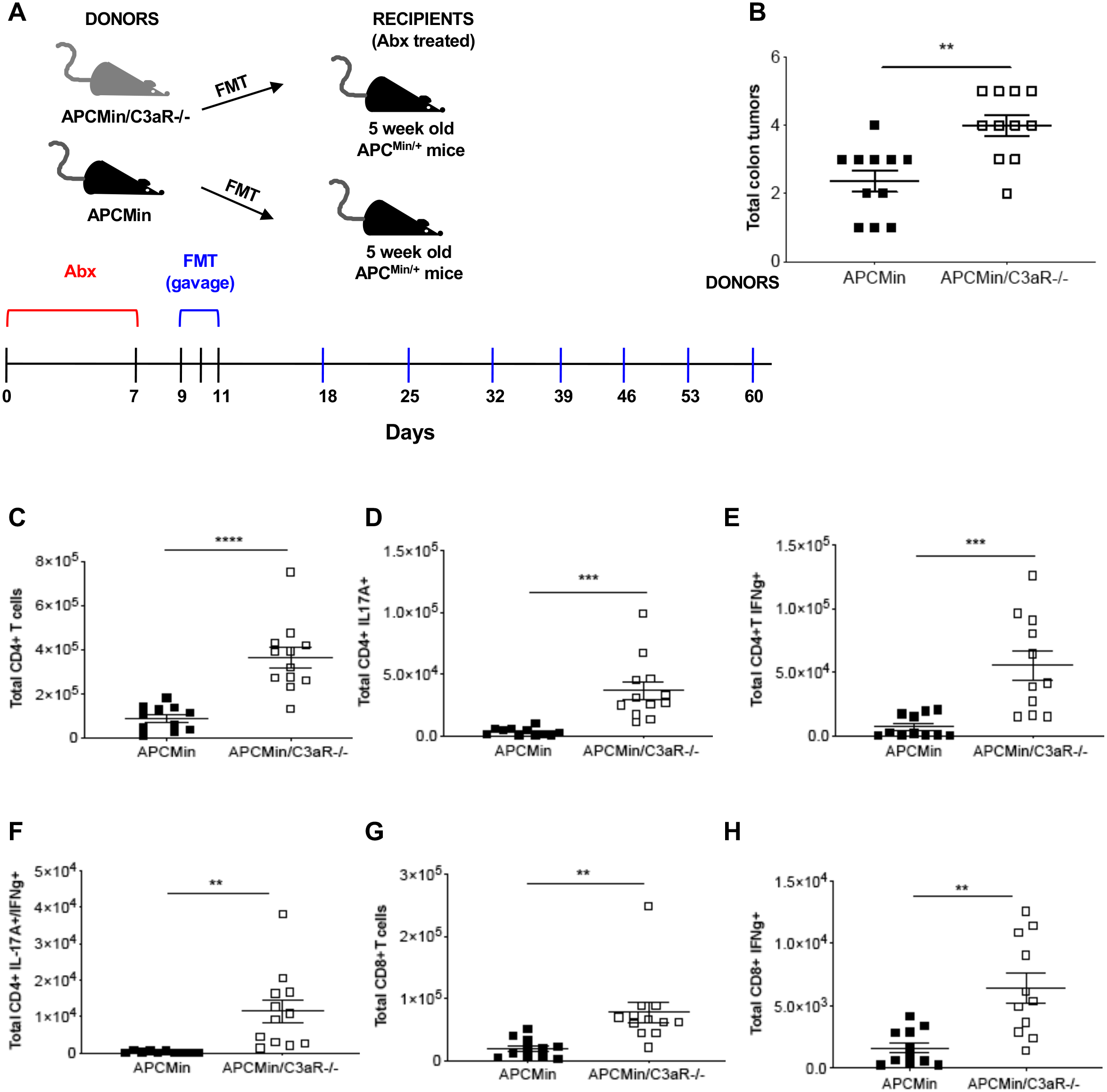
Transplantation of the APCMin/+/C3aR-/- microbiota transfers colon tumorigenesis to APCMin/+ mice. (A) 5-week-old APC^Min/+^ mice (recipients) (n=22) were treated for 1 week with broad spectrum antibiotics (abx). 48 h after antibiotics treatment, the recipient mice were transplanted via oral gavage with the gut microbiota of 12-week-old APC^Min/+^ or APC^Min/+^/C3aR-/- mice (donors) for 3 consecutive days and again once/week for 7 weeks. (B) Tumor count in the colon of recipient mice was performed at the end of the experiment. (C-H) Flow cytometry analysis of cLP infiltrating lymphocytes showing total cell number of (C) CD4+T cells, (D) Th1 cells, (E) Th17 cells, (F) Th1/17 cells, (G) CD8+ T cells and (H) Tc cells as calculated in Fig. 2. Shown are the results of two independent experiments with a minimum of 5 mice/group. Significance was calculated using unpaired t-test (** p< 0.01; *** p< 0.001; **** p<0.0001).

### Loss of C3aR affects the composition of the fecal and tumor-associated microbiota in APC^Min/+^mice

Although several lines of evidences have shown that the complement system plays a fundamental protective role against pathogens, there is a lack of studies regarding the possible role of complement in controlling a pro-tumorigenic microbiota. Recently, C5aR has been shown to play an important role in the composition of skin microbiota and the associated inflammatory immune responses (Chehoud et al., 2013). Further, C3a has been demonstrated to be a potent anti-microbial agent (Nordahl et al., 2004). To better understand whether the results obtained with the microbiota transplantation were the consequence of a tumor-associated microbiota or a change in microbiota composition driven by C3aR deficiency, we characterized the microbiota before and after tumor development. For this purpose, we collected the fecal pellet from tumor free eight-week-old and tumor-bearing 12-week-old APC^Min/+^ and APC^Min/+^/C3aR-/- mice. Bacterial DNA was extracted from fecal pellets as detailed in the experimental procedures. The V5-V6 hypervariable regions of bacterial 16S rRNA gene were amplified and the obtained libraries were sequenced on the MiSeq Illumina platform. Compared to controls, we observed higher alpha diversity in eight-week-old APC^Min/+^/C3aR-/- and C3aR-/- mice measured by observed ASV, Faith and Shannon index (Fig. 7a). The alpha diversity normalized then in 12-week-old mice, although there was still a clear separation among the two mouse strains in terms of beta diversity (Fig. 7b). These results support the concept that the fecal microbiota that precedes tumor growth is highly diversified within the same group. In contrast, upon tumor development, the selection of defined bacterial species results in reduced intra-group and enhanced inter-group diversification. When examining the differences in the most abundant phyla, we observed that, already at 8 weeks when the mice in our colony are devoid of tumors in both small intestine and colon, in APC^Min/+^/C3aR-/- mice there was increased Bacteroidetes and Proteobacteria and evident reduction of Firmicutes as compared to APC^Min/+^ mice. These differences became even more striking at 12 weeks when we could see the first differences in colon tumorigenesis (Fig. 7c). Notably, Bacteroidetes are significantly up-regulated in the stool-associated microbiota of CRC patients and have been correlated with elevated Th17 in their healthy mucosa (Ahn et al., 2013; Sobhani et al., 2011). Although the fecal microbiota reflects the disease status when comparing CRC patients with healthy controls, several lines of evidences recently showed that the microbiota specifically associated with the mucus layer and with the tumor itself differs from the fecal microbiota and might more closely reflect the changes occurring in the tumor microenvironment (Flemer et al., 2017). To understand whether the differences found in the fecal microbiota were mirrored at the tumor site, in addition to the previously described analysis, we sequenced the mucus- and tumor-associated bacteria in 12- week-old APC^Min/+^ and APC^Min/+^/C3aR-/- mice. As shown in Fig. 8a, the tumor-and mucus-associated bacterial phyla were strikingly similar between APC^Min/+^ and APC^Min/+^/C3aR-/- mice and were dominated by Firmicutes followed by Bacteroidetes. These results closely mirror the bacterial composition in CRC, where it has been shown that Firmicutes is the predominant phylum in the cancerous tissues followed by Bacteroidetes (Gao et al., 2015). It is worth noting that the bacterial composition in tumor and mucus of the two mouse strains was more similar than that in the stool. These results suggest that the physiologic and metabolic alterations occurring in the tumor microenvironment might represent the main constraint that dictates the specific composition of the microbial community.

**Figure 7.**
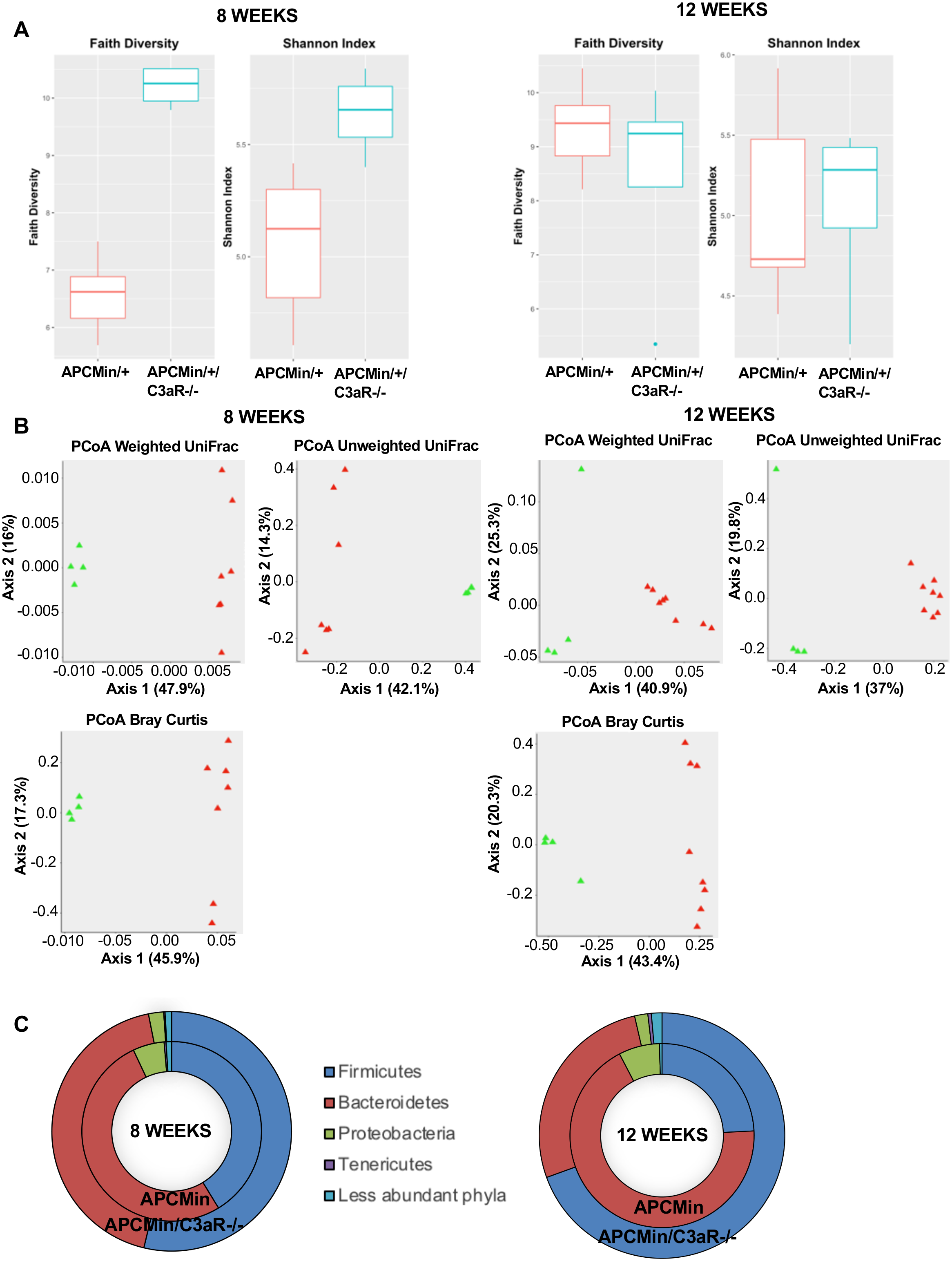
Loss of C3aR affects the composition of the fecal microbiota in APCMin/+mice. Bacterial DNA was extracted from the feces of 8-and 12-week-old APC^Min/+^ and APC^Min/+^/C3aR- /- mice, and bacterial profiling was performed by sequencing the V5-V6 hypervariable of 16S rDNA using Illumina MySeq platform. (A) Plots showing alpha diversity evaluated by observed ASV, Faith and Shannon Index in 8-and 12-week-old mice. (B) PCoA showing the beta diversity evaluated by the three inferred Beta Diversity metrics (weighted UniFrac, unweighted UniFrac and Bray-Curtis) in 8- and 12-week-old mice. (C) Doughnut charts showing phylum abundance at 8 and 12 weeks. Significance was calculated in A using Kruskal-Wallis test followed by a pairwise Wilcoxon as post-hoc test; in B using PERMANOVA test and in C using DESeq R-package. (** p<0.01)

**Figure 8.**
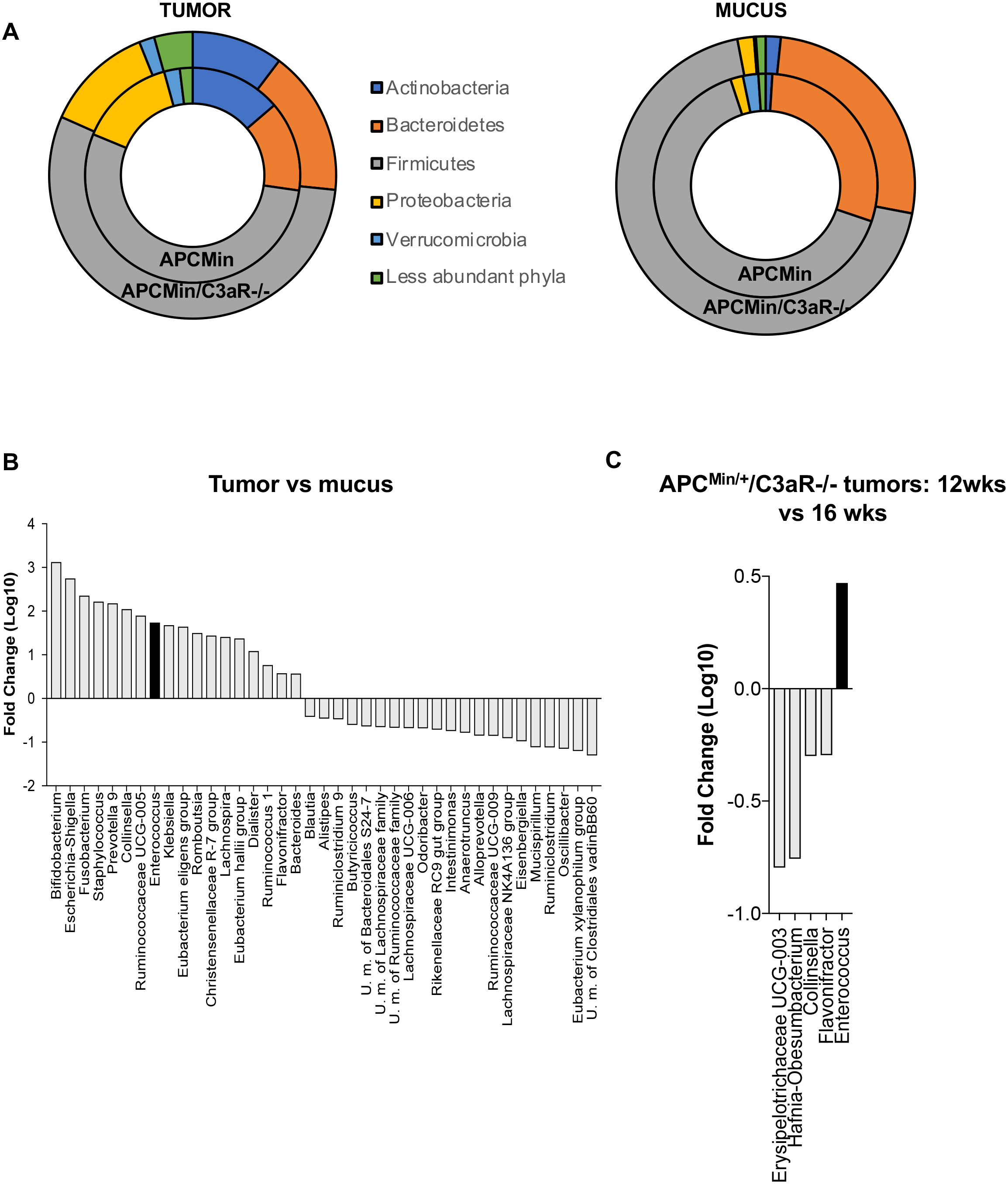
Impact of C3aR on tumor- and mucus-associated microbiota species. Bacterial DNA was extracted from colonic tumors and mucus of 12-week-old APC^Min/+^ and APC^Min/+^/C3aR-/- mice and metagenomic analysis was performed. (A) Doughnut charts showing the bacterial composition of mucus and colon of 12-week-old APC^Min/+^ and APC^Min/+^/C3aR-/- mice. (B) Bacterial species up-or down-regulated in the tumor *vs* mucus in APC^Min/+^/C3aR-/- mice. Bacterial species up-and down-regulated in tumors of 12- *vs* 16-week-old APC^Min/+^/C3aR-/- mice. Significance was calculated in A and B by DESeq R-package; in C using 1-way Anova with Bonferroni post-test.

### Colon tumors of APC^Min/+^/C3aR-/- mice harbor an *E. faecalis* strain that supports tumor growth

The results of the metagenomic analysis on tumor- and mucus-associated microbiota showed that among the most represented species in the phylum Firmicutes both in APC^Min/+^ and APC^Min/+^/C3aR-/- mice there was *E. faecalis*, which was found up-regulated in the tumors of APC^Min/+^/C3aR-/- mice compared to the mucus and increased over time in the tumor microenvironment (Fig. 8b-c). *E. faecalis* is a Gram-positive bacterium that preferentially colonizes the colonic mucosa and has been proposed as a possible CRC driver due to its ability to produce superoxide and hydrogen peroxide that induce DNA damage and support malignant transformation (Sears and Garrett, 2014; Wang et al., 2015). To understand the effect of *E. faecalis* on tumor growth, we isolated these species from the tumors of either APC^Min/+^ or APC^Min/+^/C3aR- /- mice: colon tumors were collected and homogenized as described in the experimental procedures, and serial dilutions were plated on tryptic broth (TB) agar plates. Single colonies were picked and plated on fresh agar plates in order to ensure purity and were subsequently characterized by colony PCR and 16S sequencing.

Considering that the production of pro-oxidative reactive oxygen species (ROS) is well described in *E. faecalis* and could be relevant for its pro-inflammatory activity, we evaluated C11 and C19 for their ability to produce ROS. As shown in Fig. 9a, C19 showed enhanced ROS production. Therefore, we next analyzed whether the two *E. faecalis* had a different effect on colon tumor growth. To accomplish this, five-week-old APC^Min/+^ mice were treated with broad-spectrum antibiotics for one week, then left untreated or administered with C11 or C19 as detailed in the experimental procedures and sacrificed at 12 weeks (Fig. 9c). As shown in Fig. 9c, 66.7% of the mice receiving C19 and 16.7% receiving C11 showed rectal bleeding throughout the experiment while no bleeding was observed in control mice. Accordingly, the administration of C19 resulted in significantly higher tumor number and load compared to C11 (Figure 9d-e). There were no significant differences in the tumors in the small intestine (Suppl. Fig. 5a-b). We also characterized colon infiltrating immune cells by flow cytometry and found that the administration of C19 was associated with higher infiltration by Th17 and Th1/Th17 cells in the colon lamina propria, however, without reaching statistical significance when compared to mice receiving C11 (Fig. 9f-i). In addition, as previously shown with FMT, we found no differences in immune cell infiltration in the mLN (Suppl. Fig. 5c-e). These results suggest that lack of C3aR, an essential component for anti-microbial functions, promotes the establishment of an inflammatory fecal microbiota that drives immune infiltration in the tumor microenvironment, where more virulent bacteria, such as *E. faecalis* (C19), are selected and able to persist, further boosting inflammation.

**Figure 9.**
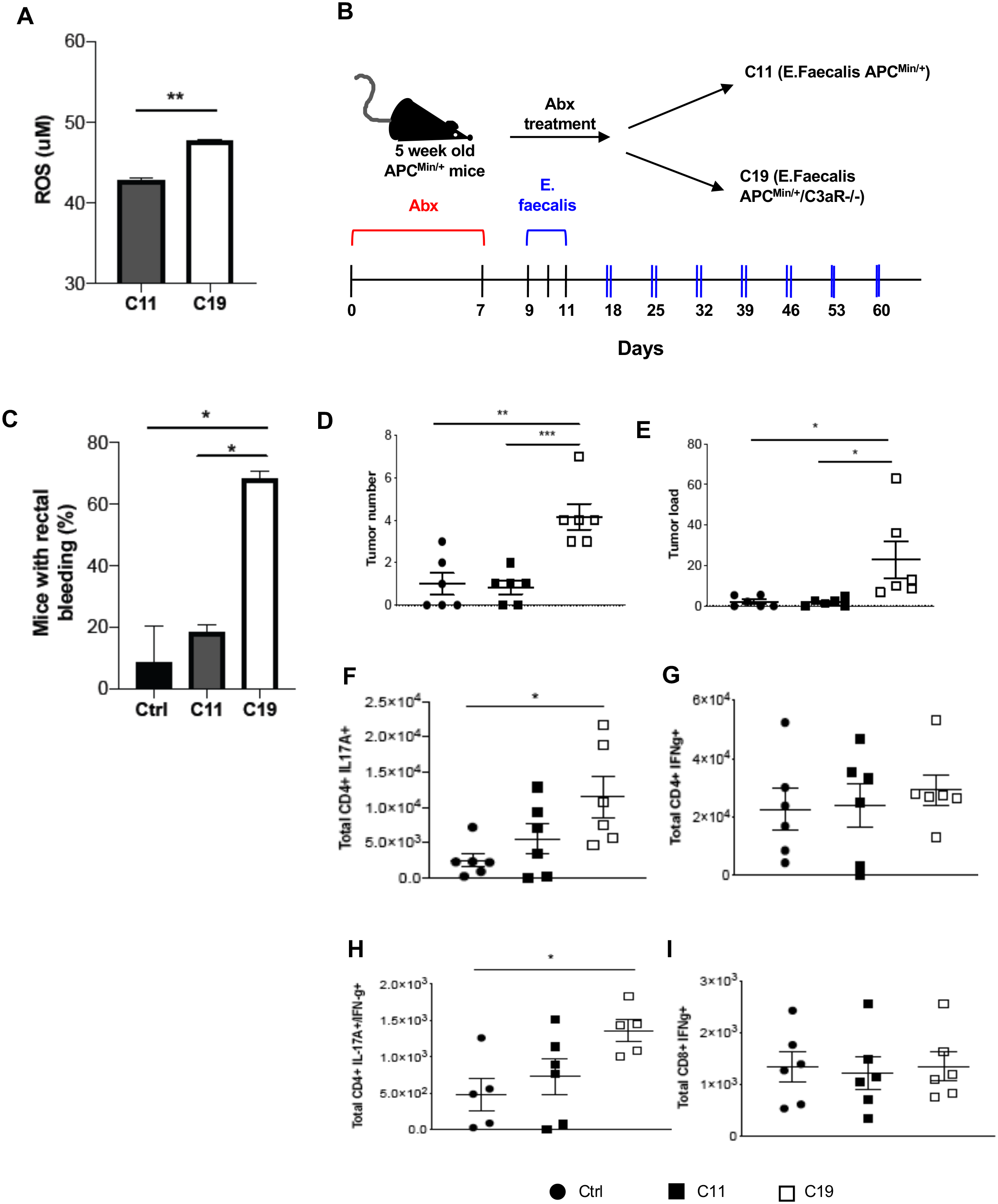
Colon tumors of APC^Min/+^/C3aR-/- mice harbor a more inflammatory *E. faecalis* strain. (A) 1.2 x 10^8^ C11 or C19 bacteria cell cultures were used to assess ROS production with the Amplex Red Hydrogen Peroxide/Peroxidase Assay Kit. Measurements were taken after 60 min of incubation. (B) 5-week-old APC^Min/+^ mice (n=6) were treated for one week with broad-spectrum antibiotics and subsequently administered with 10^9 CFU of *E. faecalis* isolated from APC^Min/+^ (C11) or APC^Min/+^/C3aR-/- (C19) mice or left untreated for 7 weeks. (C) Percentage of mice with rectal bleeding in each group. (D) Tumor number and (E) tumor load in the colon of the mice was evaluated at the end of the experiment. (F) Th1 cells, (G) Th17 cells, (H) Th1/17 cells and (I) Tc cells in the colon lamina propria of tumor-bearing mice were assessed by flow cytometry. Significance in panel A was calculated using unpaired t-test; in C-I using 1-way Anova with Bonferroni post-test (* p<0.05; ** p< 0.01; *** p< 0.001).

## DISCUSSION

The role of complement in intestinal homeostasis and CRC is not well understood.

Here we report for the first time that downregulation of the receptor for complement anaphylatoxin C3a (*c3ar1*) and CpG island methylation occur in CRC. By reverse-translating this finding we were able to recapitulate colonic tumorigenesis in APC^Min/+^, which usually show a prevalence of tumors in the small intestine (Guglietta et al., 2016; Moser et al., 1990). Therefore, by knocking down C3aR we were able to generate a reproducible mouse model that more closely mirror the tumorigenic process occurring in humans. Furthermore, we found that loss of C3aR profoundly affects the inflammatory infiltrate of colonic tumors and results in a pro-inflammatory fecal and tumor-associated microbiota. Inactivation of the central component of the complement cascade C3 was shown to exacerbate chronic intestinal inflammation via upregulation of inflammatory cytokines, demonstrating that activation of the complement cascade is a key regulator of intestinal homeostasis (Elvington et al., 2015). In one of the few prior studies investigating the role of complement in CRC, the authors showed that the complement anaphylatoxin C5aR exacerbates tumor development in an AOM/DSS model of inflammation-driven CRC, by recruiting IL-1β producing neutrophils (Ning et al., 2015). Significant modulation of the complement system has been also reported in cancer molecular subtype 2 (CMS2) and CMS4 human CRC (Guinney et al., 2015). In addition, a very recent study reported that up-regulation of the complement regulatory protein CD55, which inhibits the central complement component, C3 convertase, correlates with decreased disease-free survival in human CRC (Olcina et al., 2018).

Several mechanisms could potentially mediate C3aR down-regulation or inactivation in human CRC. In this context it is important to note that, while genetic mutations in the complement system as a group occur at a significant higher rate than in any other gene for several cancers, most individual complement genes mutate at a lower rate than many canonical oncogenes and oncosuppressors (Ding et al., 2010; Olcina et al., 2018). Accordingly, we found that in CRC patients as compared to healthy controls there is a significantly higher frequency of C3aR methylation occurring in those regions that are likely to result in mRNA expression changes. Therefore, methylation may represent the main cause of C3aR downregulation in human CRC. Mechanistically we found that lack of C3aR resulted in significant accumulation of Th17 and Th1/Th17 cells in the colon lamina propria and in the tumors. The role of IL17 in CRC is highly debated. Indeed, production of IL-17A has been shown to be important for preserving the integrity of the epithelial barrier (Kumar et al., 2016; Lee et al., 2015). However, data in patients and mouse models of CRC showed that IL17 production by different cell sources promotes exacerbation of the inflammatory process and fosters cancer cell proliferation (Goktuna et al., 2016; Grivennikov et al., 2012; Wang et al., 2014). Furthermore, a recent study by Omenetti and collaborators showed that Th17 cells can be defined by their ability to produce high levels of inflammatory cytokines and by the activation of inflammation-related pathways (Omenetti et al., 2019). Our flow cytometry and RNASeq data clearly show that, besides their pronounced ability to produce IL-17A, the cells enriched in the colon lamina propria and tumors of APC^Min/+^/C3aR-/- mice produced significantly higher amount of IFN-*γ* as compared to their counterpart in APC^Min/+^ mice. In tumors and colon lamina propria of APC^Min/+^/C3aR-/- mice, we consistently found up-regulation of several signal transducers and activators of transcription, such as STAT-1 and STAT-4, and increased levels of inflammatory cytokines, such as TNF-α, IL-22 and IL-1β. The induction of mucosal immune responses is largely dependent on the host microbiota and several reports highlighted the effect of innate immune mechanisms in modifying the gut flora (Fulde et al., 2018; Salzman et al., 2010; Vaishnava et al., 2011). Besides being regarded as an important link between innate and adaptive immune responses, the complement system is fundamentally a first line of defense against pathogens. Moreover, C3a, the ligand of C3aR, has been demonstrated to exert anti-bacterial functions (Nordahl et al., 2004). Despite this evidence, there are no studies in the literature evaluating the effect of C3aR on the gut microbiota. However, Chehoud and collaborators found that genetic ablation and antagonism of C5aR correlated with an inflammatory microbiota in the skin, suggesting that complement anaphylatoxins receptors may also play a role in shaping the gut flora (Chehoud et al., 2013). Consistent with data in literature showing that C3aR-/- mice are unable to efficiently clear Gram-negative bacteria, we found that the fecal microbiota of APC^Min/+^/C3aR-/- and C3aR-/- mice was characterized by higher abundance of Gram-negative bacteria such as Bacteroidetes and Proteobacteria (Hollmann et al., 2008; Kildsgaard et al., 2000). This preceded tumorigenesis and was further enhanced during tumor development. The fecal microbiota established in the absence of C3aR was likely the main driver of Th1, Th17 and Th1/Th17 cells in the colon of APC^Min/+^ mice that received fecal microbiota transplantation from APC^Min/+^/C3aR-/- mice. Indeed, the tumor- and mucus-associated microbiota were strikingly similar between APC^Min/+^ and APC^Min/+^/C3aR-/- mice. Nevertheless, we report the identification of a strains of *E. faecalis*, in the tumors of APC^Min/+^/C3aR-/- mice, that displayed more pathogenic features compared to the same species isolated from APC^Min/+^ tumors. A few studies in humans reported significantly higher levels of *E. faecalis* in CRC patients compared to healthy controls, however the mechanisms linking *E. faecalis* to CRC remain unclear (Balamurugan et al., 2008; Wang et al., 2012). In our work, when used in single bacteria transfer experiments in vivo, *E. faecalis* from APC^Min/+^/C3aR-/- mice resulted in higher tumor load and number compared to *E. faecalis* from APC^Min/+^ mice. Mechanistically, the small but significant increase in ROS production observed in *E. faecalis* from APC^Min/+^/C3aR-/- mice could explain the increased tumorigenic potential, therefore confirming previous finding (Huycke et al., 2002; Huycke and Moore, 2002). Based on our findings regarding the role of C3aR in regulating the microbiome and immune infiltrate during tumor development, it is tempting to speculate that similar mechanisms may be at play in other human cancers and especially in those arising in surfaces exposed to the external environment where innate immune defense mechanisms and microbiome play an important role (Gomes et al., 2014; Janakiram and Rao, 2014; Maru et al., 2014; Senol et al., 2014).

Future studies will be needed to assess the mechanisms responsible for *c3ar1* methylation and to determine whether C3aR downregulation could be used as biomarker or exploited for therapeutic purposes.

## METHODS

### Animals

C57BL/6J-*ApcMin*/J (referred to as APC^Min/+^) and C57BL/6J (referred to as WT) mice were bred and maintained in our SPF (Specific Pathogen Free) animal facility at European Institute of Oncology and at the Medical University of South Carolina. C3aR-/- mice on C57BL/6J background were a kind gift from Dr. Bao Lu and Dr. Carl Atkinson. APC^Min/+^/C3aR-/- mice were generated by crossing C57BL/6J-*ApcMin*/J with C3aR-/- mice in our animal facilities. All animal experiments were performed in accordance with the guidelines established in the Principles of Laboratory Animal Care (directive 86/609/EEC) and approved by Italian Ministry of Health and under approved protocols by the Institutional Animal Care and Use Committee at MUSC (IACUC- 2017-00165; IACUC-2020-01022)

### AOM/DSS

Twelve-week-old WT and C3aR-/- mice were intraperitoneally injected with 10 mg/Kg of azoxymethane (AOM; Sigma-Aldrich). After 7 days, mice were administered with 1.5% (w/v) DSS (TdB Consultancy) in their drinking water for 1 week followed by 2 weeks of recovery. DSS/recovery cycles were repeated three times as previously described (Neufert et al., 2007). Mice were weighed every other day. At the end of the experiment, mice were sacrificed, tumors in the colon were counted and their diameter measured using a sliding caliper. Tumor load for each mouse was calculated by adding the average diameter of all tumors.

### Spontaneous colonic tumorigenesis and FACS analysis of infiltrating immune cells

APC^Min/+^ and APC^Min/+^/C3aR-/- mice were sacrificed at different time points starting at the age of 5 weeks until 28 weeks and tumors in the colon were counted. Colon, tumors and mesenteric lymph nodes (mLN) were used to characterize the immune cells by FACS. Single cell suspensions were prepared from mLN following standard protocols. Colon lamina propria cells were isolated as previously described with minor modifications (Lefrancois and Lycke, 2001).

For isolation of tumor-associated immune cells, colon polyps were shaken in PBS, 1% BSA, 10 mM EDTA in order to remove epithelial cells, and further digested for 30 minutes at 37°C with Collagenase VIII (Sigma-Aldrich) in complete medium with shaking.

For flow cytometry staining, cells were incubated with anti-FcR antibody (clone 24G2) and stained with the following surface antibodies: anti-CD45.2 (clone 104, eBioscience), Ly6G (clone 1A8), CD3 (clone 17A2, eBioscience), Ly6C (clone AL-21), CD11b (clone M1/70), CD4 (cloneRM4- 5), CD8a (53-6-7), B220 (clone RA3-6B2), I-A/I-E (clone M5/114.15.2), CD11c (clone HL3), F4/80 (MB8, eBioscience), CD103 (clone 2E7), CD25 (clone PC61), anti-IFN-*γ* (clone XMG1.2), anti-IL-17A (clone TC11-18H10), anti-FoxP3 (clone FJK-16s eBioscience). For cytokine staining cells were permeabilized with Cytofix/Cytoperm buffer (BD) according to manufacturer instructions. For Foxp3 staining the permeabilization was performed using the FoxP3 permeabilization buffer (eBioscience). All antibodies were purchased from BD Pharmingen unless otherwise specified. Samples were acquired with FACSCanto II or Fortessa LSR (BD Bioscience) and analyzed with FlowJo software (TreeStar).

### Fecal microbiota transplantation and gavage with E. faecalis

For fecal microbiota transplantation (FMT), fecal material was obtained from 12-week-old APC^Min/+^ and APC^Min/+^/C3aR-/- donor mice. Briefly, fecal pellet and cecal contents were harvested from the mice, dissolved in sterile PBS at 100 mg/ml, passed through a 70-μm strainer in order to eliminated insoluble debris, and frozen in Columbia broth with 20% glycerol. At the time of administration, aliquots were thawed, centrifuged to eliminate the glycerol-containing medium and resuspended at 100 mg/ml for gavage. This procedure allowed us to use the same material throughout the experiment, thereby avoiding confounding effects due to fecal and mucus material coming from different animals. Five-week-old APC^Min/+^ mice were used as FMT recipient. Recipient mice were treated for one week with the broad-spectrum antibiotics in the drinking water, rested for 48 h with water without antibiotics, followed by gavage with 200 μl /mouse of the previously prepared fecal and mucus material for 3 consecutive days in the first week and once a week for the next 6 weeks. For *E. faecalis* administration, we followed a treatment scheme similar to the one described for FMT. *E. faecalis* isolated from APC^Min/+^ and APC^Min/+^/C3aR-/- mice were grown in aerobic condition at 37°C with terrific broth (TB). CFUs were determined along the growth phase from serial dilution plated on TB plates at 37°C for 24 h. Next, single colonies were grown overnight in TB broth, restarted at OD_600_ of 0.05 in fresh TB and grown to OD_600_ of 0.8. 10^9^ CFU from each bacteria culture were administered via oral gavage to each mouse and after the first 3 consecutive administrations, we performed 2 gavages/week for the remaining 6 weeks. At the end of the treatment animals were sacrificed, colon tumors counted, and single cell suspensions were prepared from colon and mLN.

### Fecal, mucus and tumor microbiota profiling

Fecal pellets were harvested from 8- and 12-week-old APC^Min^ and APC^Min/+^/C3aR-/- mice and used to extract DNA with G’NOME DNA isolation kit (MP) following a published protocol (Furet et al., 2009). V5-V6 hypervariable regions of bacterial 16S rRNA gene were amplified and processed with a modified version of the Nextera protocol (Manzari et al., 2014). The obtained metabarcoding libraries were sequenced by using the MiSeq Illumina platform with a 2×250 paired end (PE) approach. Metagenomic amplicons were analyzed by applying the BioMaS pipeline (Fosso et al., 2015): (i) The paired-end reads were merged into consensus sequences using Flash (Magoc and Salzberg, 2011) and subsequently dereplicated as previously described (Edgar, 2010) maintaining the consensus sequence; (ii) The remaining non overlapping PE reads were considered for further analysis only if after the low-quality region trimming (Phred quality cut-off = 25) both read ends were ≥50 bp long; (Mariathasan et al.) Both the merged sequences and the unmerged reads were matched against the RDP (Ribosomal Database Project) database (release 10.29)(Cole et al., 2009) by Bowtie2 (Langmead and Salzberg, 2012). The mapping data were filtered according to two parameters: identity percentage (≥97%) and query coverage (≥70%); (iv) Finally, all mapped reads fulfilling the settled filters were taxonomically annotated using the Tango tool (Alonso-Alemany et al., 2014). Assigned genera were filtered considering as present only the ones for which at least five reads per samples were present. The read counts were normalized using an approach similar to the RPKM (Reads per kilo-base per million): normalized count = assigned reads / (total assigned reads at the rank level/1,000,000). Significant differences at the genus and species level were calculated with the DESeq R-package (Anders and Huber, 2010). A tree representing the phylogenetic relationship between the Amplicon Sequence Variants (ASV) was produced by using the QIIME1 package (Caporaso et al., 2010). The rarefaction curves were inferred and plotted by using an in house developed R script relying on the rarefy function of the vegan package. Alpha and Beta diversity analysis were performed using the phyloseq R (McMurdie and Holmes, 2013). In particular, the observed ASVs, the Shannon and the Faith Indexes were used as richness and alpha diversity measures. The weighted and unweighted UniFrac and the Bray-Curtis metrics were used for the Beta diversity inference (Chang et al., 2011). Difference in the inferred alpha diversity indices were measured by using the Kruskal-Wallis test followed by a pairwise Wilcoxon as post-hoc test (p-values corrected by using the Benjamin-Hochberg (BH) procedure). The PERMANOVA test was used to compare groups in beta diversity data, using 999 permutations.

To isolate bacterial DNA from mucus scraped from the colon we used the same protocol used to isolate bacterial DNA from the feces. For the isolation of tumor-associated bacteria, tumors were homogenized in sterile phosphate buffered saline solution (PBS), centrifuged, and both supernatants and pellets were subjected to depletion of host DNA by using the QIAamp DNA Microbiome kit (Qiagen) before proceeding with bacterial DNA extraction with the same protocol described for feces.

### Colony PCR

Bacteria associated to mucus and feces were plated on agar plates and screening of different colonies was performed using colony PCR. Briefly, cells from a single colony were picked from agar plates by using a sterile pipette tip and resuspended into lysis buffer (Tris/EDTA, 0.2 % SDS, and 10 mM EDTA). The lysate was then centrifuged, and the supernatant was collected and used as PCR template. PCR amplification was performed on the 16SrRNA amplifying V1-V4 with 28F (5’-GAGTTTGATCNTGGCTCAG-3’), 907R (5’-CCGTCAATTCMTTTRAGTTT-3’) and V5-V7 with 1392R (5′-ACGGGCGGTGTGTRC-3′), 926F (5′-AAACTYAAAKGAATTGACGG-3′). PCRs were performed with Phusion High-Fidelity DNA Polymerase (2 U/µL) (New England Biolabs) in a total volume of 50 μl. Each cycle consisted of 60 s at 98 °C, 45 s at 63 °C, and 60 s at 72 °C, with a final extension of 5 min. The PCR products were subjected to 1 % agarose gel electrophoresis for analysis and purified with QIAquick PCR Purification Kit (Qiagen) according to the manufacturer’s protocol.

### Reactive oxygen species (ROS) production by *E. faecalis*

*E. faecalis* isolated from APC^Min/+^ and APC^Min/+^/C3aR-/- mice were grown in aerobic conditions at 37°C in TB. Single colonies of both *E. faecalis* were grown until the stationary phase. As template for ROS detection, the culture was centrifuged at 13.000 rpm for 2 minutes and the supernatants were filtered and collected. Alternatively, to start the reaction, 1.2 x 10^8^ bacteria cell cultures (bacteria cells plus supernatant) were added to the plate. The Amplex Red Hydrogen Peroxide/Peroxidase Assay Kit (Invitrogen) was used to detect ROS production following manufacturer’s instructions. All the measurements were performed in 96-well plates, using GloMax® Discover Microplate Reader (Promega) equipped for excitation at 540 nm and detection at 590. Measurements were taken after 60 minutes of incubation.

### RNA-Seq analysis of colon and colonic tumors

For RNASeq analysis, 12-week-old APC^Min/+^, WT, C3aR-/- and APC^Min/+^/C3aR-/- mice (4 mice/group) were sacrificed. The proximal (first 3 cm adjacent to the caecum) and distal colon (the final 3.5 cm adjacent to the rectum) and the single tumors were immediately preserved in 2 ml of RNAlater stabilization solution (Qiagen) according to manufacturer instructions and stored at −20°C. For RNA extraction, tumors and colons were homogenized in Trizol (500 ul/50 mg weight), centrifuged at 13000 rpm, the supernatant was loaded on Qiagen Mini Kit columns and treated as indicated in the manufacturer instructions. RNA concentration was quantified by Nanodrop and quality and integrity were evaluated by using a bioanalyzer (Agilent Technologies). Only samples with a RIN>8 were used for downstream applications.

Briefly, 1μg of RNA was used to prepare cDNA with the TruSeq RNA kit following the recommendations for low sample preparation protocol (Illumina). Samples were sequenced on an Illumina MiSeq instrument at a depth of 35×10^6 reads per sample. Reads in FASTQ format were quantified at the gene level using featureCounts and the count table was delivered to edgeR for the differential expression analysis using the GLM functionality (Liao et al., 2014; McCarthy et al., 2012). GO analysis was conducted to identify DEG at the biologically functional level. The identified DEGs were uploaded to the online software Advaita Bioinformatics, which was used to integrate functional genomic annotations.

A false discovery rate (FDR) cutoff of 5% and minimum fold change of 1.5 (log2FC=0.6) was applied to determine differential expression.

RNASeq data are deposited on the ArrayExpress website (accession number: E-MTAB-8500).

### Analysis of C3aR methylation and mutations in patients with CRC

Data on C3aR expression were obtained from the R2 Genomics Analysis and Visualization Platform (http://r2.amc.nl) or TCGA database. C3aR expression data were plotted with PRISM (GraphPad Software). Data on C3aR methylation were obtained from the human pan-cancer methylation database (Huang et al., 2015).

### Statistical analysis

Data were analyzed for normal distribution before performing statistical analyses. Values are presented as means ± standard error mean (Overman et al.) of multiple individual experiments, each carried out at least in triplicate, or as means ± SEM of replicates in a representative experiment. Comparison between two groups was determined by the Student’s t test or Mann-Whitney test. Comparison of multiple groups was carried out by one-way or two-way ANOVA followed by Bonferroni post-test correction using GraphPad Prism software version 8 as indicated in the figure legends. All statistical tests were two-sided, and *P*<0.05 was considered statistically significant, unless otherwise specified.

## COMPETING INTERESTS

The authors have declared that no conflict of interest exists.

## ACKNOWLEDGMENTS

The authors wish to acknowledge Roger Johnson for manuscript editing; Dr. Carl Atkinson and Dr. Stephen Tomlinson for providing essential reagents and mice; Dr. Luca Rotta at the IEO Genomics Unit for assistance with RNA sequencing; Dr. Graziano Pesole for assistance with microbiota sequencing; Dr. Sara Carloni for help with culturing bacterial strains; American Cancer Society-Institutional Research Grant, AIRC and Fondazione Umberto Veronesi to S.G.

## AUTHOR CONTRIBUTION

S.G. conceived the study and the experimental setup, performed and analyzed experiments and wrote the manuscript with input from all authors; C.K. performed experiments and analysis and wrote the manuscript; L.M.W., M.D.R. and G.H. performed analysis of RNASeq data; B.F. and M.M. performed microbiota sequencing and analysis; E.M. assisted with mouse experiments; S.E.A. gave input for microbiota experiments and analysis.

## Notes

https://www.ebi.ac.uk/arrayexpress

### Competing Interest Statement

The authors have declared no competing interest.

