## Supplemental material for "Loss of C3aR induces immune infiltration and inflammatory microbiota in a new spontaneous model of colon cancer"

#### SUPPLEMENTAL INFORMATION

**Supplementary Figure 1. Loss of C3aR expression does not significantly impact the functional and phenotypic composition of mLN-infiltrating lymphocytes.**

Single cell suspensions from mLN of $APC^{Min/+}$ and $APC^{Min/+}$/C3aR-/- mice were analyzed by FACS and total number of (A) CD4+T cells, (B) Th17 cells, (C) Th1 cells, (D) Th1/Th17 cells, (E) CD8+T cells, (F) Tc cells were obtained. A minimum of 5 animals/group was used and significance was calculated using unpaired t-test (** $p< 0.01$; ****$p>0.0001$).

**Supplementary Figure 2. Differential gene expression analysis in distal colon of $APC^{Min/+}$, $APC^{Min/+}$/C3aR-/-, C3aR-/- and WT mice.**

(A-F) Summary plot showing the number of significant differentially expressed genes between distal colons of (A) C3aR-/- *vs*. WT mice (n=4); (B) $APC^{Min/+}$ *vs*. WT mice; (C) $APC^{Min/+}$/C3aR-/- *vs*. C3aR-/- mice; (D) $APC^{Min/+}$/C3aR-/- *vs.* WT mice; (E) $APC^{Min/+}$ *vs.* C3aR-/- mice and (F) $APC^{Min/+}$/C3aR-/- *vs* $APC^{Min/+}$. The plots display log fold change against log counts per million for each gene with the red points representing significant differentially expressed genes, and the horizontal blue lines indicating the threshold. Significance was calculated using the edgeR function decideTestsDGE with default p-value adjustment method (Benjamini-Hochberg false discovery rate, FDR set at 0.05, and a custom fold change threshold of 2 (log2FC = 1).

**Supplementary Figure 3. Transplantation of the $APC^{Min/+}$/C3aR-/- microbiota does not affect the number of tumors in the small intestine (SI) and the functional profile of mLN-infiltrating lymphocytes.**

5-week-old $APC^{Min/+}$ mice (recipients) (n=22) were treated for one week with ampicillin. 48 h after antibiotics treatment, the recipient mice were transplanted via oral gavage with the gut microbiota of 12-week-old $APC^{Min/+}$ or $APC^{Min/+}$/C3aR-/- mice (donors) for 3 consecutive days and again once/week for 7 weeks. (A) Total tumor count in the SI and (B) in the different portions of the SI: duodenum (D), jejunum (J) and ileum (I) of recipient mice was performed at the end of the experiment. (C-H) Flow cytometry analysis of mLN infiltrating lymphocytes showing in (C) total CD4+T cells, (D) Th17 cells, (E) Th1 cells, (F) Th1/Th17 cells, (G) CD8+ T cells and (H) Tc cells. Shown are results of 2 independent experiments with a minimum of 5 mice/group. Significance was calculated in A and B using 1-way Anova and 2-way Anova respectively with Bonferroni post-test and in C-H using unpaired T test (** $p < 0.01$; *** $p < 0.001$; **** $p < 0.0001$).

**Supplementary Figure 4. E. faecalis from $APC^{Min/+}$/C3aR-/- tumors does not affect small intestinal tumors.**

(A) Total tumor number in the SI and (B) in each compartment of the small intestine of $APC^{Min/+}$ mice treated with C11 or C19 strains of *E. faecalis*. (C) Th17 cells, (D) Th1 cells, (E) Tc cells in the mLN of treated mice were assessed by flow cytometry. Significance was calculated in panels a-d using 1-way Anova with Bonferroni post-test.

### Supplementary Figure 1

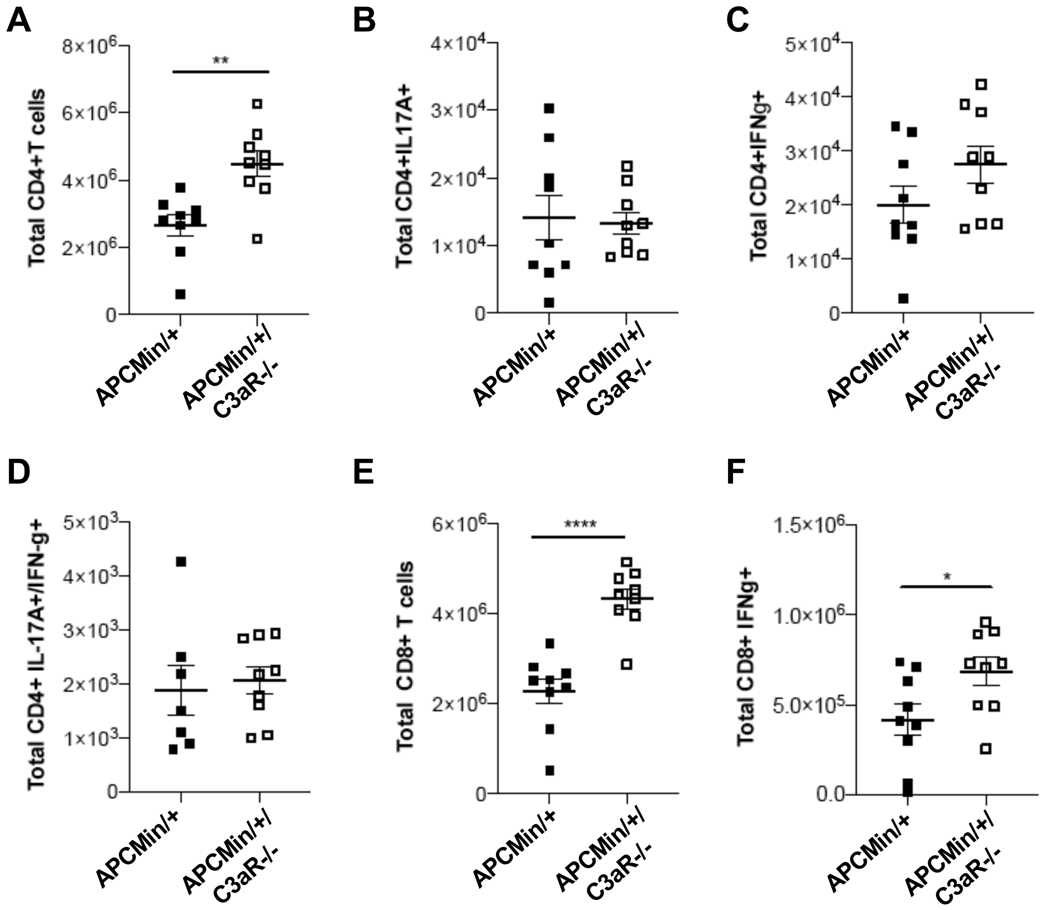

### Supplementary Figure 2

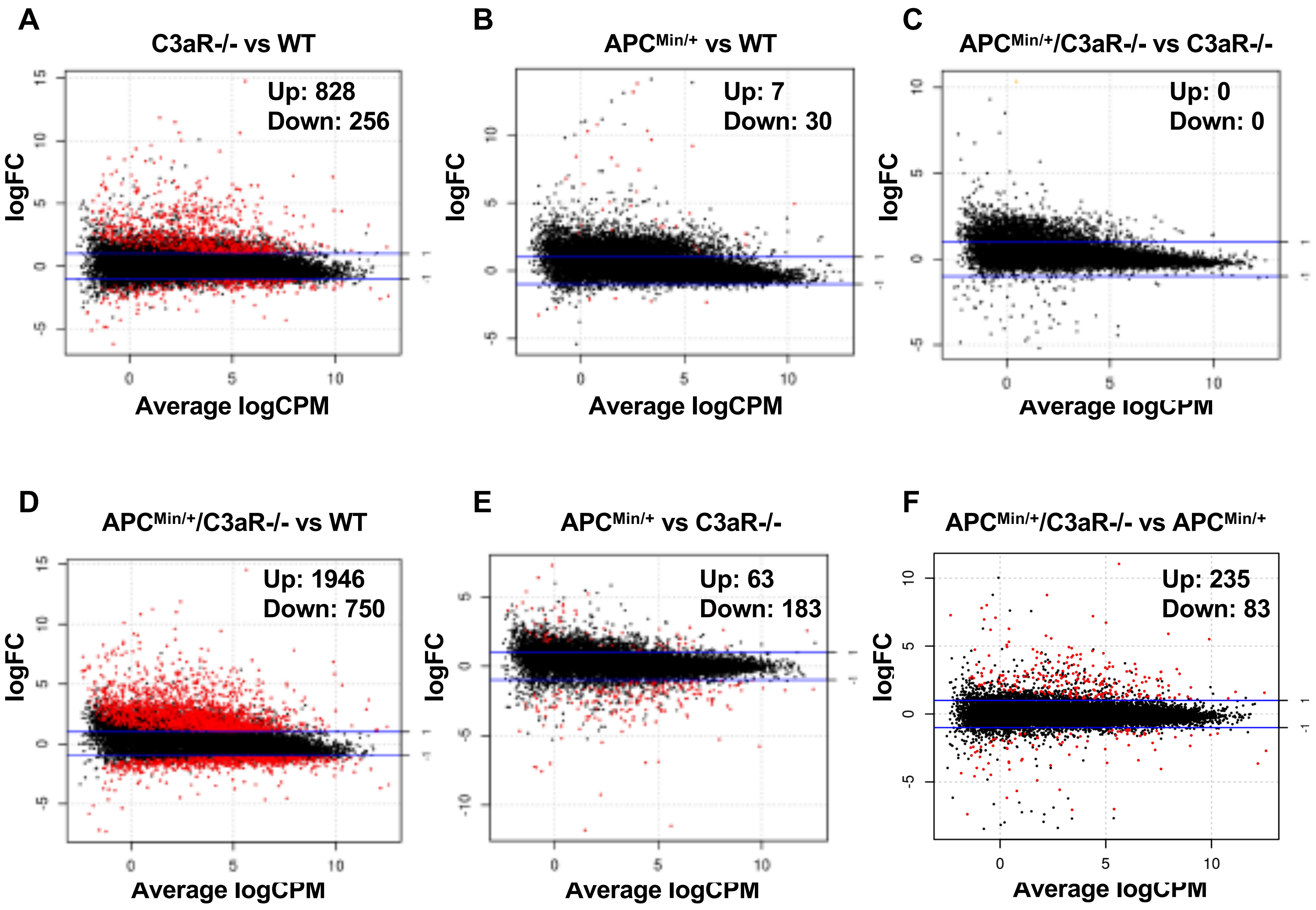

### Supplementary Figure 3

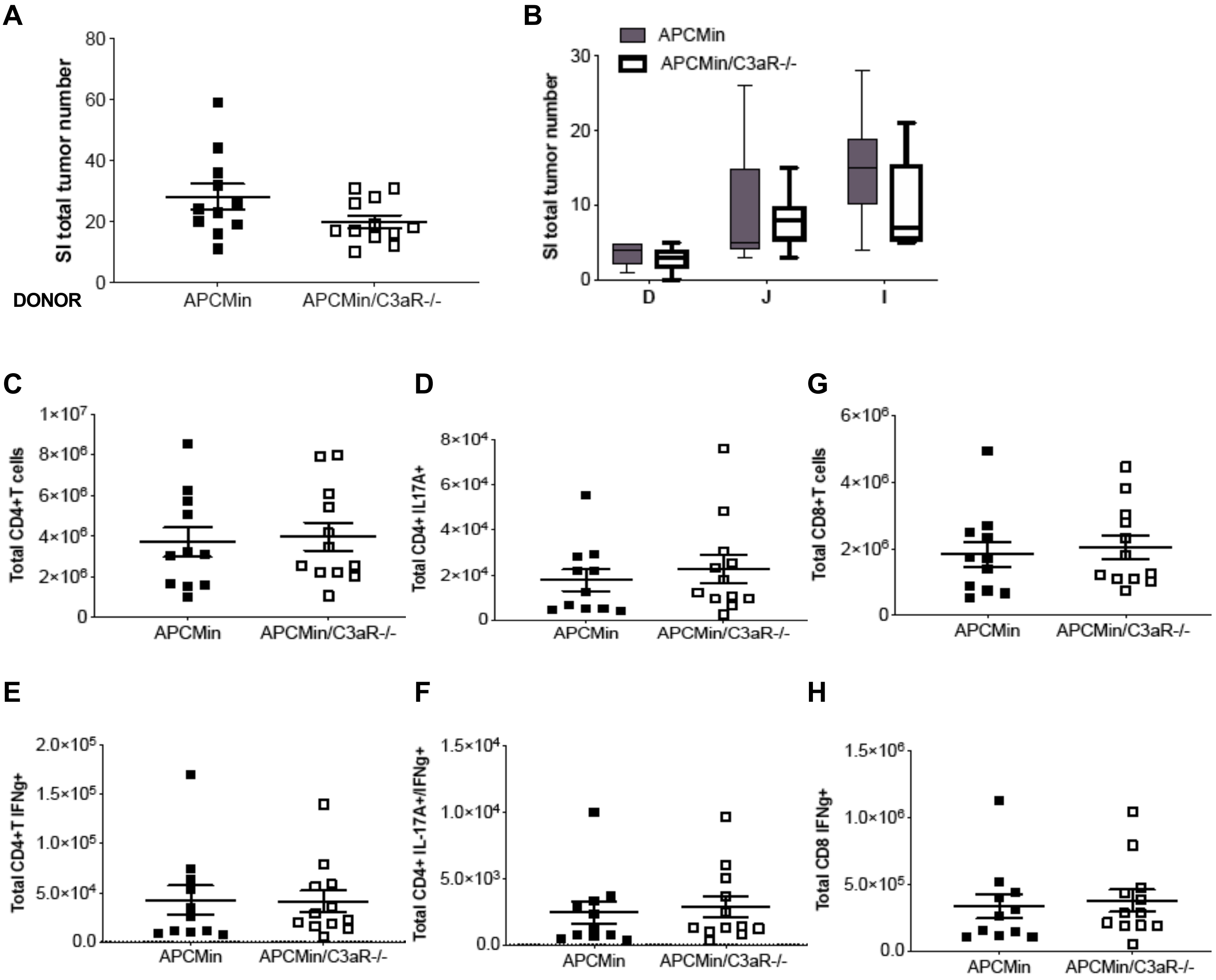

#### Supplementary Figure 4

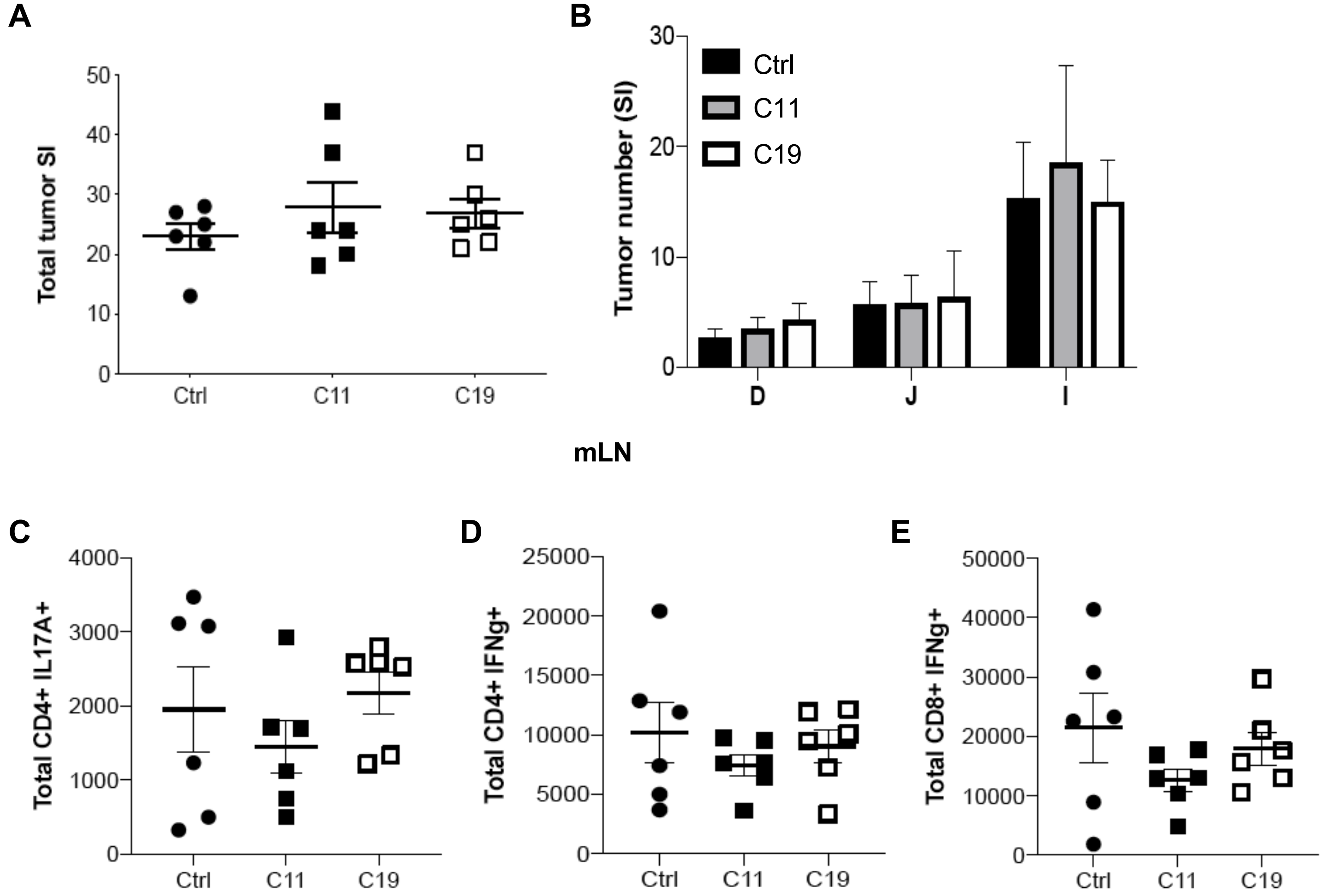
